# Investigating the conformational dynamics of SARS-CoV-2 NSP6 protein with emphasis on non-transmembrane 91-112 & 231-290 regions

**DOI:** 10.1101/2021.07.06.451329

**Authors:** Amit Kumar, Prateek Kumar, Kumar Udit Saumya, Rajanish Giri

## Abstract

The NSP6 protein of SARS-CoV-2 is a transmembrane protein, with some regions lying outside the membrane. Besides, a brief role of NSP6 in autophagosome formation, this is not studied significantly. Also, there is no structural information available till date. Based on the prediction by TMHMM server for transmembrane prediction, it is found that the N-terminal residues (1-11), middle region residues (91-112) and C-terminal residues (231-290) lies outside the membrane. Molecular Dynamics (MD) simulations showed that NSP6 consisting of helical structures, whereas membrane outside lying region (91-112) showed partial helicity, which further used as model and obtain disordered type conformation after 1.5 microsecond. Whereas, the residues 231-290 has both helical and beta sheet conformations in its structure model. A 200ns simulations resulted in the loss of beta sheet structures, while helical regions remained intact. Further, we have characterized the residue 91-112 by using reductionist approaches. The NSP6 (91-112) was found disordered like in isolation, which gain helical conformation in different biological mimic environmental conditions. These studies can be helpful to study NSP6 (91-112) interactions with host proteins, where different protein conformation might play significant role. The present study adds up more information about NSP6 protein aspect, which could be exploited for its host protein interaction and pathogenesis.

**Graphical Abstract:** 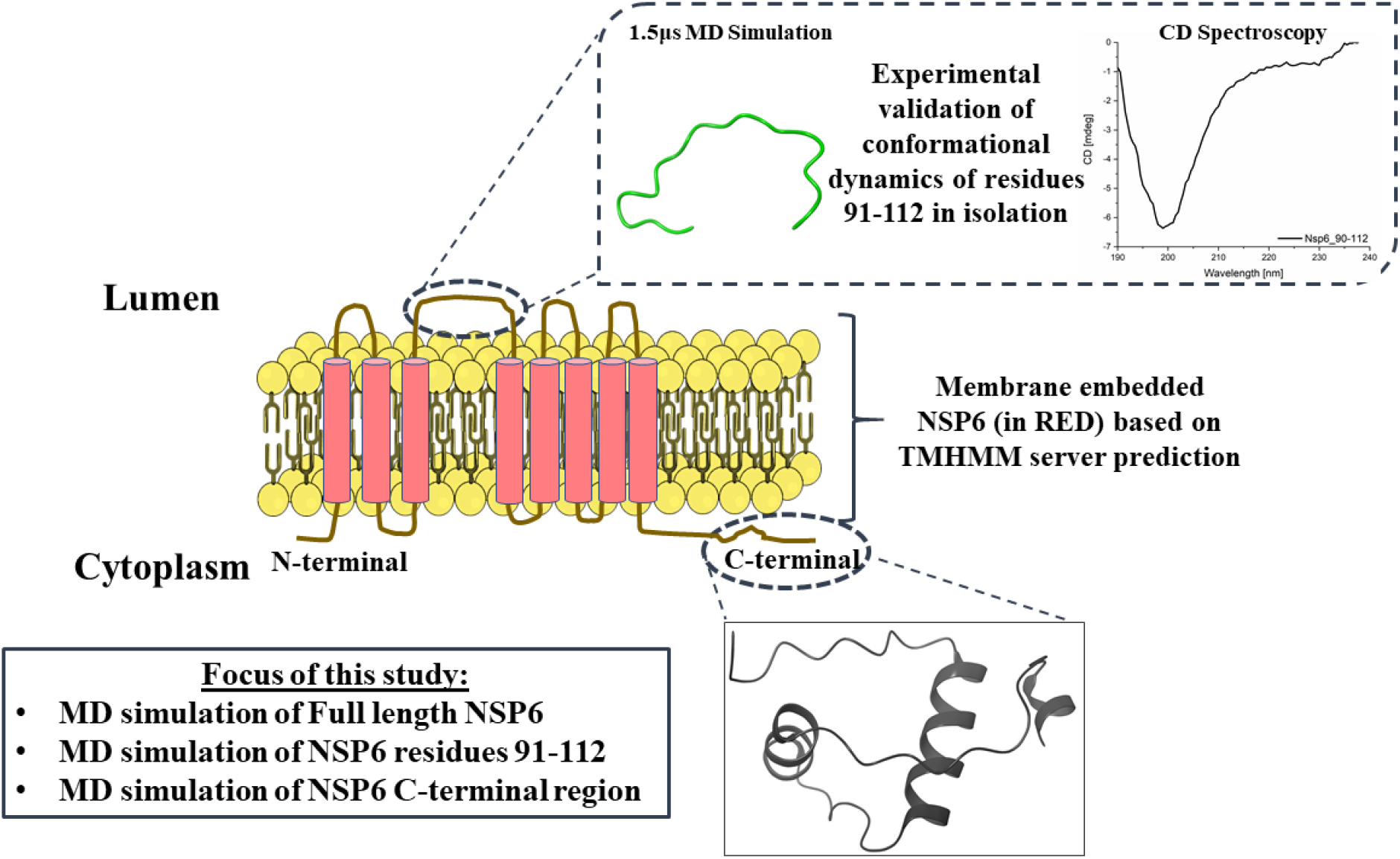

The schematic representation of NSP6 membrane topology and conformational dynamics of residue 91-112. The N-terminal and C-terminal are shown in cytoplasmic side based on the experimental evidence on coronaviruses reported by Oostra et al., 2008. The membrane anchoring domain are shown based on the TMHMM server prediction.

## Introduction

Severe acute respiratory syndrome-coronavirus 2 (SARS-CoV-2) has positive-sense single-stranded RNA genomes□(∼□30 kilobases), which encodes sixteen nonstructural proteins (1). There are numerous articles which described in details about the SARS-CoV-2 life cycle, genome, proteome, and functioning mechanisms (2–5). Among non-structural proteins (NSPs), very less is known about NSP6, specifically there is no 3D structure is available till date. Gordon et al., stated that NSP6 interact with host proteins, although the mechanism still remains elusive (4). A yeast two hybrid assays shows that NSP6 interact with other non-structural viral proteins of SARS-CoV (6). The information on NSP6 protein is very limited such as structural studies, host protein interactions and many other, which are still needed to address. It is reported that The NSP6 protein is a six-pass membrane spanning protein facing it’s both terminal at cytosolic side (7–9). The NSP6 protein along with NSP3 and NSP4 is found to form double membrane vesicle (10).

The coronaviruses (CoV-1 and CoV-2) NSP6 protein interacts with sigma receptor 1 (*SIGMAR1*) (4). Further, it has been reported for CoV-1 NSP6 that it plays role in autophagy, even in the absence of its C-terminal region (8). Considering the NSP6 interaction with host proteins, the structural characterization of either full length or partial NSP6 domain is necessary to understand the SARS-CoV-2 pathogenicity mechanism. However, the surrounding environment play important role in the conformational dynamics of protein or its interacting regions (11). The change in conformation is governed by the covalent (i.e., phosphorylation, ubiquitination) and/or non-covalent events i.e., binding of ions, lipids, drugs, proteins etc. and environmental influences such as macromolecular crowding, osmolyte, pH, and temperature (11–16). Factors influencing the change in proteins conformational dynamics ultimately decide the fate of subsequent protein signaling event (11, 17).

In the present study, we have used TMHMM server to predict the regions lying outside the membrane for their potential to interact with host proteins. Based on computational modelling and simulation, we have characterized NSP6 full length, membrane outside lying region of NSP6 and NSP6-C terminal region (CTR). We experimentally investigated the synthetic peptide of membrane outside faced region (residues 91-112) and studied in isolation for its conformational dynamics to validate our predicted results. The short peptide sequence give invaluable information regarding their conformation in terms of their capabilities to regulate the cellular process or in context of complete global structure (18, 19). Our studies found that this region is disordered like in isolation and gain helical conformation in the presence of TFE and SDS, suggest its propensity to gain structure while interaction with particular partners.

## Material and methods

### Chemicals and reagents

The NSP6 peptide (residues 91–112) “-NH2-VMRIMTWLDMVDTSLSGFKLKD-COOH-”, with purity >88% was purchased from Gene script, USA. Organic solvents such as Trifluoroethanol (TFE) with ≥99% purity was purchased from Sigma-Aldrich. Lyophilized peptide was dissolved in 10 mM sodium phosphate buffer at a concentration of 10 µM.

### Molecular Dynamics (MD) Simulation

We have utilized I-TASSER, PepFold, and RaptorX (20–22) web-servers for constructing the 3D models for the NSP6 full-length (NSP6-FL), NSP6 (residues 91-112) and NSP6-CTR, respectively. The resultant models were then prepared using Chimera by adding missing hydrogens and missing sidechains were completed in residues (23). We have utilized Charmm36m forcefield in Gromacs v5 on high performance cluster (HPC) of IIT Mandi, where simulation setup was built by placing the protein structure (NSP6 91-112 and NSP6-CTR) in a cubic box with a distance of 10Å from each edge along with SPC water model and 0.15M NaCl salt concentration. NSP6 is a membrane bound protein, therefore for full length NSP6 model lipid membrane environment was provided. The simulation setup for NSP6 full-length was built using CHARMM-GUI web server, where 250 molecules of neutrally charged lipid, Dioleoylphosphatidylcholine (DOPC) (24). After solvation, all systems were charge neutralized with counterions. The steepest descent method was used to attain an energy minimized simulation system until the system was converged within 1000 kJ/mol. Further, the equilibration of the system was done to optimize solvent in the environment. Using NVT and NPT ensembles within periodic boundary conditions for 100ps each, the system was equilibrated. The average temperature at 300K and pressure at 1 bar were maintained using Nose-Hoover and Parrinello-Rahman coupling methods during the simulation. All bond-related constraints were solved using SHAKE algorithm. The final production run was performed for 100ns, 1500ns (1.5 microsecond; µs), and 200ns, for NSP6-FL, NSP6 (91-112), NSP6-CTR, respectively.

All trajectory analysis calculations and visualizations were performed using Chimera, maestro, vmd and Gromacs command for calculating the helicity, root mean square deviation (RMSD), root mean square fluctuation (RMSF), and radius of gyration (Rg) for protein structure compactness.

### Liposome preparation

The liposomes were prepared, as described earlier (15). Briefly, the neutral lipid DOPC (dioleoyl-phosphatidyl-ethanolamine) was purchased from Avanti Polar Lipids (Alabaster, AL). The chloroform from the lipid solution was removed using a rotary evaporator at 40°C, and the dry lipid films were hydrated in 50 mM phosphate buffer (pH 7.4). The final concentration of the DOPS liposomes was 24.69 mM, respectively. The resulting suspension was freeze-thaw-vortex in liquid nitrogen and water at 60 °C, following which the lipids were extruded 25 times through the mini extruder (Avanti Polar Lipids, Inc. USA) through cut off filter of 100nm polycarbonate membrane to prepare uniform Large unilamellar vesicles (LUV).

### Circular Dichroism spectroscopy

JASCO machine (Jasco J-1500 CD spectrometer) was used for CD data recording. 5 μM peptide sample were prepared in 10 mM phosphate buffer, pH 7.0. The peptide was kept in organic solvents (TFE) with increasing concentration from 0 to 50%, and far-UV (190–240 nm) spectra were recorded in 1 mm quartz cuvette. Similarly, the peptide (10 µM) was assessed for structural changes in Sodium Dodecyl Sulfate (SDS) and liposome DOPS. All the spectra were recorded at a scan speed of 50 nm/min with a response time of 1s and 1 nm bandwidth and three technical repeats. The equivalent spectra of buffers were recorded and subtracted from the spectra of the test samples. Further, the smoothing of CD spectra was done by Savitsky-Golay fitting at 5 points of smoothing window and second polynomial order.

### Fluorescence spectroscopy

We monitored the intrinsic Trp fluorescence intensity in NSP6 (90-112). A 5μM peptide in 10mM sodium phosphate buffer (pH 7.0) was prepared with increasing TFE and SDS concentration. Emission spectra were recorded from 300 to 500 nm at 295 nm excitation wavelength in a Horiba Fluorolog-3 spectrofluorometer. The individual negative blank was subtracted from each test sample (14).

## Results and discussion

### TMHMM server predicted NSP6 (residues 91-112) lies outside the membrane

These outcomes make us to look into the residues 91-112, which lies outside the membrane and might play role in interaction with host protein or other unpredicted functions. Therefore, our aim was to characterize this region in particular (91-112). Firstly, the TMHMM server was used to characterize the amino acid of NSP6 for their transmembrane region, inside and outside of membrane region **(Figure 1)**. Similarly, Benvenuto et al., showed 7 transmembrane region in NSP6 of coronaviruses by using TMHMM server and investigated the effect of mutation in NSPs and orf10 in their adjacent region (7).

**Figure 1:**
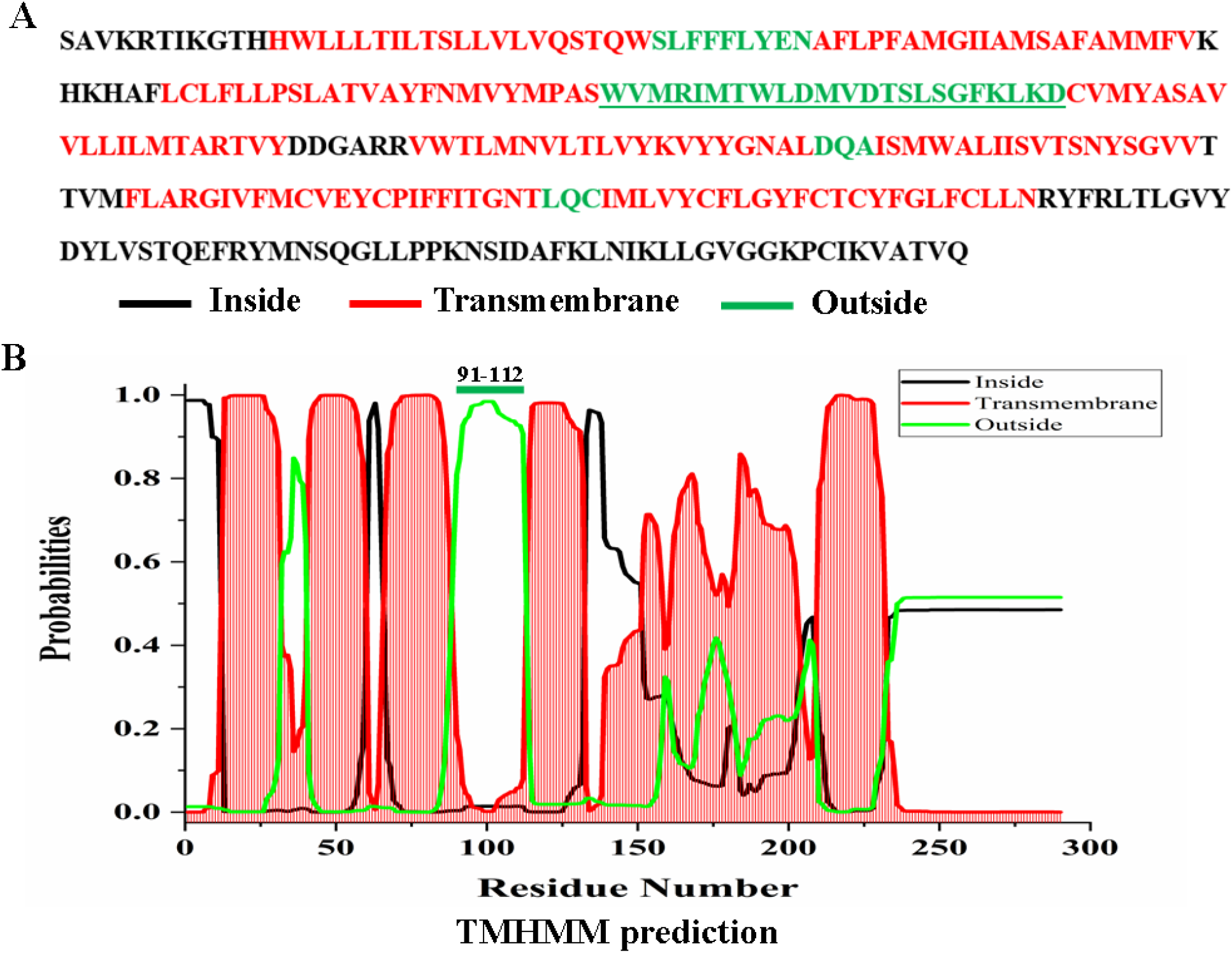
**(A)** The primary sequence of SARS CoV-2 NSP6 full length with color pattern showing the inside, outside and transmembrane region. **(B)** TMHMM server prediction with color pattern showing the residue 91-112 lies outside the membrane.

### MD simulations of full length NSP6, NSP6 (91-112) and NSP-CTR

We have built the 3D model of full length NSP6, NSP6 C-terminal region (CTR) and NSP6 (residues 91-112). Firstly, the model built using different web-servers was simulated for appropriate simulation time. As shown in **Figure 2**, the NSP6-FL comprised multiple helical regions which were predicted to be transmembrane region (shown in red; **Figure 2A**), outside membrane regions (shown in green; residues 91-112; **Figure 2B)** and inside membrane region (shown in black; residues 231-290; **Figure 2C**). After 100 ns of MD simulations of NSP6-FL, the membrane passing helical regions were intact while the regions lying outside of the membrane showed fewer changes in secondary structure (secondary structure timeline is shown in **supplementary figure 1**). In isolation, the structure model of residues 91-112 consisted nearly 65% of helix in its structure but has shown huge structural changes after simulations. The NSP6 (residues 91-112) lost its helicity after 1.5 µs, which showed its disorder character in isolation (**Figure 2B**). This is also confirmed from secondary structure timeline shown in **supplementary figure 2**. Interestingly, the C-terminal or cytosolic region (residues 231-290) has predicted to comprise both helical and beta sheet (_243_DYL_245_ and _283_CIK_285_) conformations in its structure model built using RaptorX. Upon simulating it for 200 ns, the beta sheet structures of the NSP6-CTR region has converted to loop regions. Additionally, the helical regions were intact upto 200ns (secondary structure timeline is shown in **supplementary figure 3**).

**Figure 2:**
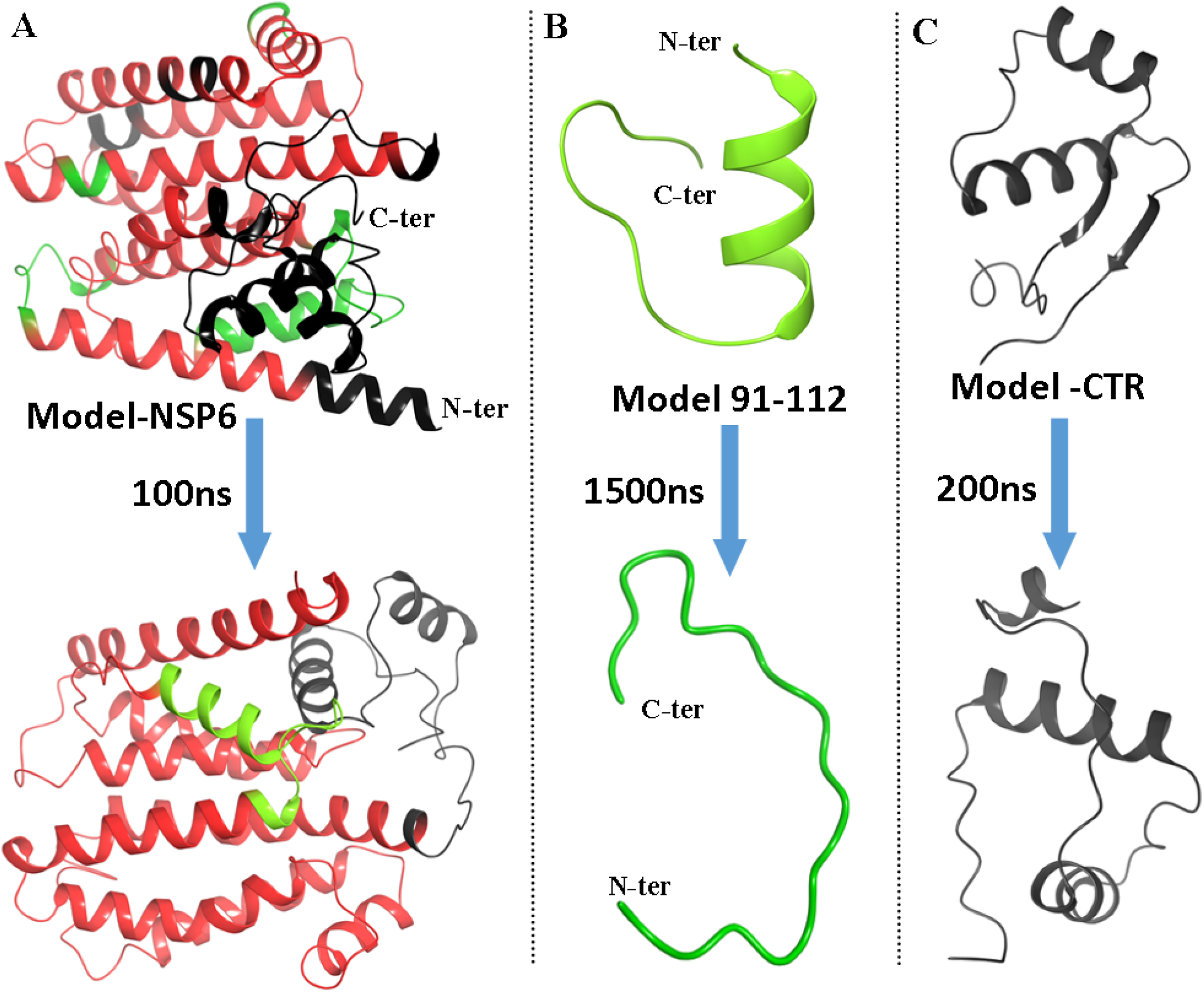
(**A**) Model of the full length NSP6 build from I-TASSER server as no 3D structure available till date and simulated for 100ns. (**B**) The NSP6 residues 91-112 in isolation showed random coil after simulation for 1.5 µs. (**C**) The NSP6-CTR residues 230-290 helical conformation after 200ns. The color pattern in protein models showing the inside (Black), outside (Green) and transmembrane region (Red).

Further, we have analyzed the time-dependent simulation frame analyses through Root Mean Square Deviation (RMSD), Radius of gyration (Rg), and Root Mean Square Fluctuation (RMSF) values (**Figure 3**). First, the modeled structure of NSP6 was simulated in the presence of membrane to check its conformation, and its RMSD values were deviating initially (up to 0.35nm till 10ns approx.) and then stabilized at 0.35nm thereafter up-to complete simulation time period. These trends were also reflected in the Rg time-dependent parameters, where values were fluctuating up to 40ns and stabilized thereafter. According to RMSF trend, the N-terminals residues fluctuate heavily. The middle regions and residue nearby to CTR also showed fluctuation (**Figure 3B**). The average helicity calculation for 100 ns simulation has shown intact helix with >90% for majority of the helices except small helices which have shown minor fluctuations (**Figure 3D**).

**Figure 3:**
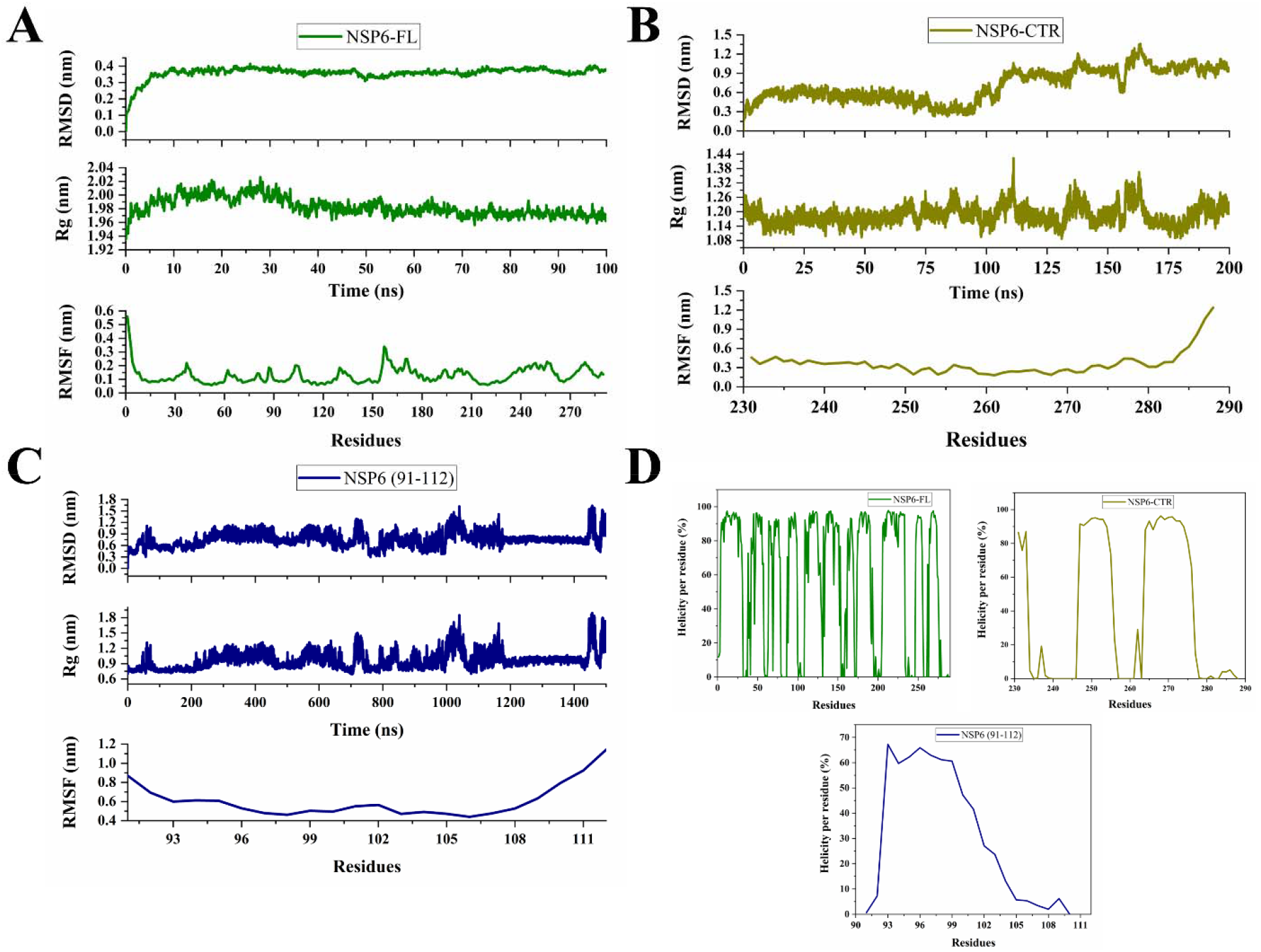
Molecular Dynamics Simulation analysis based on Root Mean Square Deviation (RMSD), Radius of gyration (Rg), and Root Mean Square Fluctuations (RMSF) of Model NSP6 full-length **(A)**, NSP6 (residues 91-112) (**B)**, NSP6-CTR region (residues 231-290) **(C)**. In figure (**D)**, the helicity percentage over throughout simulation period is shown for all three simulated structures.

Next, the inside membrane region of NSP6 (NSP6-CTR) has shown a stable trend in RMSD and Rg in initial 90 ns with an approx. average RMSD of 0.45 nm and Rg of 1.18 nm (**Figure 3B**). Afterwards, the fluctuations were increased in mean distances which may be due to change in structure of NSP6-CTR, particularly, due to transition of beta sheets to coil. The average RMSF of NSP6-CTR is in favorable range with minimal fluctuations in some residues.

The modeled structure of NSP6 (91-112) was simulated in water as it lies outer surface as per predictions. The RMSD, Rg values were found deviating upto 1200ns and stabilized thereafter. According to RMSF trend, except the residues 96-99 and residues 102-105, other residues fluctuate heavily (**Figure 3C**). The change in structural regions is also evident from **Figure 3D**, where the helicity has declined from nearly 70% for residues 93-102. This claim is further investigated by synthesized peptide based spectroscopic studies.

### NSP6 residues 91-112 intrinsically disordered region in isolation and obtain helicity in the presence of membrane mimic environment and organic solvent

Circular dichroism spectroscopy was used to monitor the conformational dynamics of NSP6 (91-112). In physiological pH and buffer conditions the peptide showed strong negative ellipticity at 198 nm, which is a characteristics of random coil conformations **(Figure 4A)**. Interestingly, in the presence of organic solvent (TFE) and SDS, peptide showed negative ellipticity at 208nm and 222 nm, which showed gain in helical conformation of peptide **(Figure 4B,C)**. The results showed that surrounding environment have strong role in conformational dynamics of NSP6 90-112 region. Organic solvent and SDS are well acknowledged for their hydrophobic and biological membrane mimic properties, respectively (14, 15, 25). Previously, these conditions are well exploited to study the numbers of protein and peptide, under the influence of hydrophobic and membrane mimic environment.

**Figure 4:**
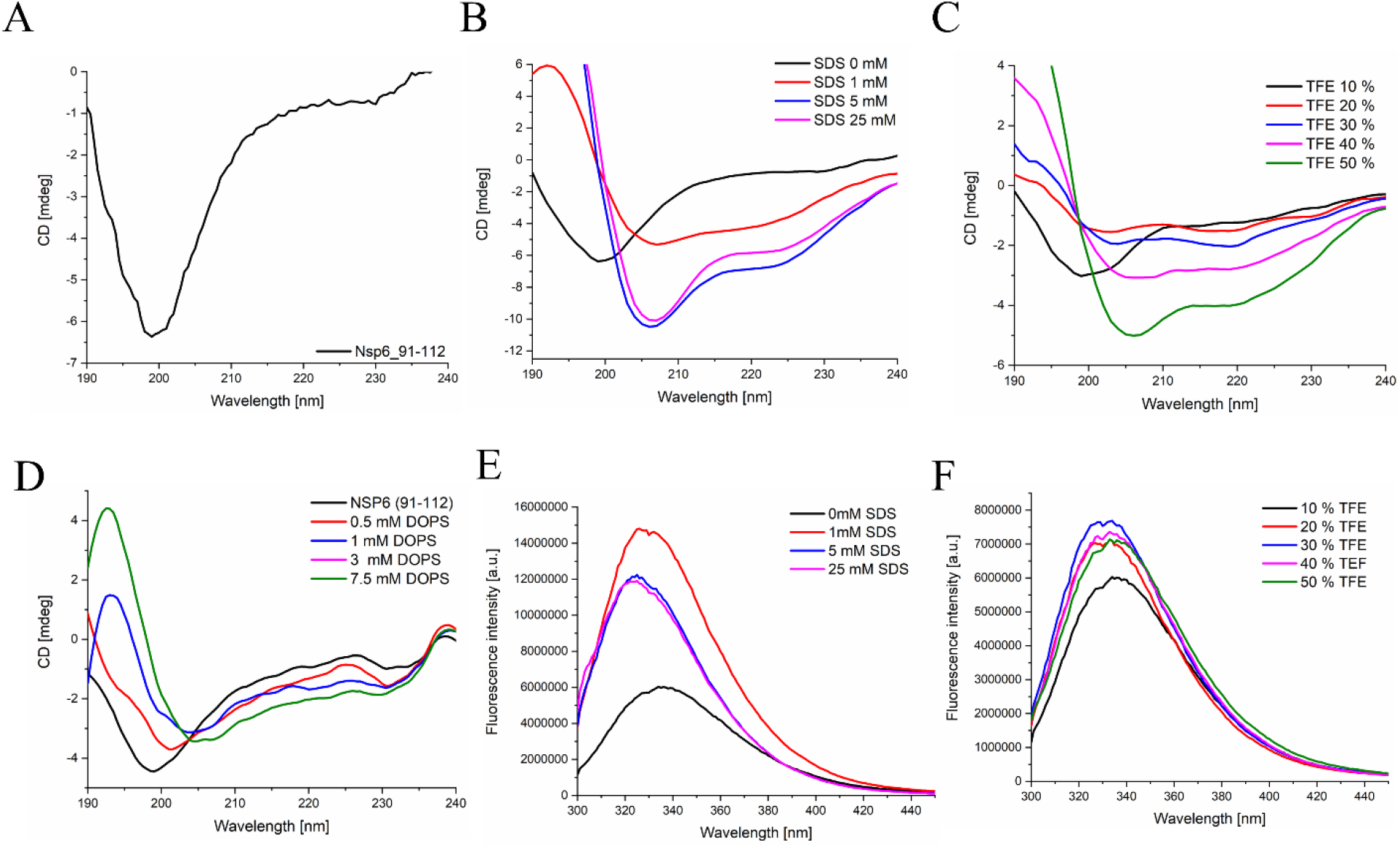
Conformation of NSP6 (91-112) studied with CD and fluorescence spectroscopy. (A) negative ellipticity of peptide under physiological buffer conditions. Effect of SDS **(B)** and TFE **(C)** showed gain in helical conformation at 222nm. Tertiary structural changes were observed with florescence spectroscopy in the presence of SDS **(D)** and TFE **(E)** showed significant blue shift under the influence of hydrophobic environment.

Furthermore, we have used the intrinsic tryptophan present in this peptide as a fluorescence probe **(Figure 4 E,F)**. Trp in the presence of non-polar/hydrophobic environment give rise to significant blue shift (25). We have observed that in the presence of TFE and SDS peptide showed blue shift, which again confirm the tertiary structural changes are happening to the peptide in these conditions and in accordance with the results obtained from the CD spectroscopy.

### Temperature induces contraction in NSP6 91-112 peptide

Further, we have characterized the peptide for its structural rigidity and changes over wide range of temperature conditions. Higher temperature conditions lead to the structural changes in the peptide, which can be relate with the gain in helicity at 222nm **(Figure 5)**. At higher temperature a phenomenon well described earlier known as contraction, which again represent the hydrophobic forces was responsible for the structural changes (15, 26, 27).

**Figure 5:**
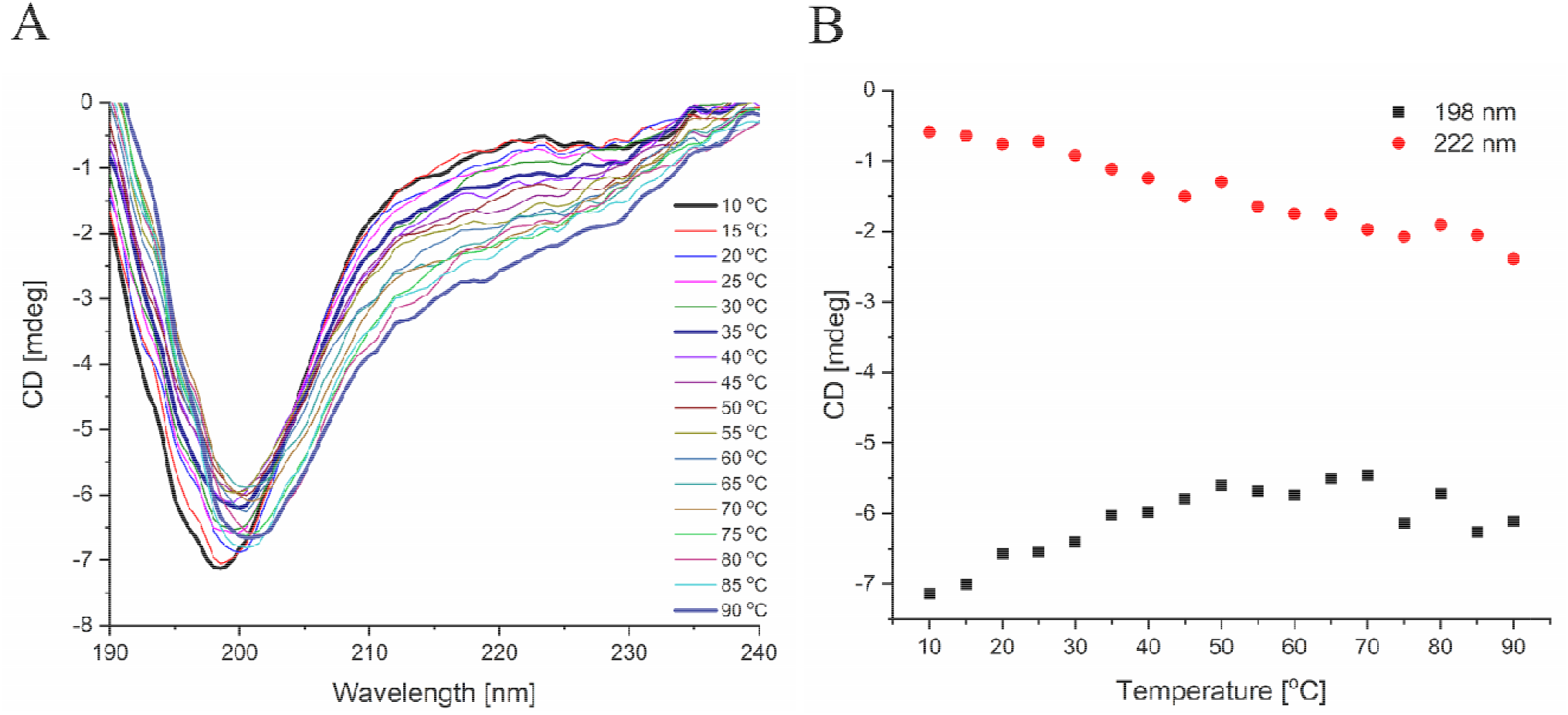
Temperature induced structural changes in NSP6 (90-112). **(A)** At higher temperature negative ellipticity was observed at 222nm shows gain in helical conformation. **(B)** Negative ellipticity at 198nm and 222nm shows change in peptide conformation.

## Conclusion

The protein structure determination by X-ray crystallography, NMR or Cryo-EM gives advantage to understand the protein in aqueous solution, which is very close to its physiological conformations. However, certain limitations still exist for certain class of protein to be studied through these high-end techniques. Similarly, the 3D structure of NSP6 is not determined till date by any of these techniques. Moreover, the SARS-CoV-2 is emerging with new variants in a short span of time, thus, the computer-aided protein structure determination is pivotal to understand the NSP6 protein conformation and for potential drug screening. The disordered/flexible conformational dynamics of non-transmembrane regions of NSP6 protein are in great importance to understand their role in protein-protein interaction and subsequent signaling events.

## Credit authorship contribution statement

RG: Conception, study design and supervision. AK, PK and KUS conducted the experiment. AK and PK acquisition and interpretation of computational data. AK, PK, and RG contributed to paper writing.

## Acknowledgments

All the authors would like to thank IIT Mandi for providing experimental and HPC facilities and Faculty Research Grant, SBS, IIT Mandi to RG. RG is thankful of DBT, Government of India (BT/11/IYBA/2018/06). AK was supported by DBT, Government of India (BT/11/IYBA/2018/06). KUS is grateful to ICMR for SRF funding.

## References

1. Giri, R., T. Bhardwaj, M. Shegane, B.R. Gehi, P. Kumar, K. Gadhave, C.J. Oldfield, and V.N. Uversky. 2020. Understanding COVID-19 via comparative analysis of dark proteomes of SARS-CoV-2, human SARS and bat SARS-like coronaviruses. Cell. Mol. Life Sci.

2. Naqvi, A.A.T., K. Fatima, T. Mohammad, U. Fatima, I.K. Singh, A. Singh, S.M. Atif, G. Hariprasad, G.M. Hasan, and Md.I. Hassan. 2020. Insights into SARS-CoV-2 genome, structure, evolution, pathogenesis and therapies: Structural genomics approach. Biochim. Biophys. Acta Mol. Basis Dis. 1866:165878.

3. Bojkova, D., K. Klann, B. Koch, M. Widera, D. Krause, S. Ciesek, J. Cinatl, and C. Münch. 2020. Proteomics of SARS-CoV-2-infected host cells reveals therapy targets. Nature583:469–472.

4. Gordon, D.E., J. Hiatt, M. Bouhaddou, V.V. Rezelj, S. Ulferts, H. Braberg, A.S. Jureka, K. Obernier, J.Z. Guo, J. Batra, R.M. Kaake, A.R. Weckstein, T.W. Owens, M. Gupta, S. Pourmal, E.W. Titus, M. Cakir, M. Soucheray, M. McGregor, Z. Cakir, G. Jang, M.J. O’Meara, T.A. Tummino, Z. Zhang, H. Foussard, A. Rojc, Y. Zhou, D. Kuchenov, R. Hüttenhain, J. Xu, M. Eckhardt, D.L. Swaney, J.M. Fabius, M. Ummadi, B. Tutuncuoglu, U. Rathore, M. Modak, P. Haas, K.M. Haas, Z.Z.C. Naing, E.H. Pulido, Y. Shi, I. Barrio-Hernandez, D. Memon, E. Petsalaki, A. Dunham, M.C. Marrero, D. Burke, C. Koh, T. Vallet, J.A. Silvas, C.M. Azumaya, C. Billesbølle, A.F. Brilot, M.G. Campbell, A. Diallo, M.S. Dickinson, D. Diwanji, N. Herrera, N. Hoppe, H.T. Kratochvil, Y. Liu, G.E. Merz, M. Moritz, H.C. Nguyen, C. Nowotny, C. Puchades, A.N. Rizo, U. Schulze-Gahmen, A.M. Smith, M. Sun, I.D. Young, J. Zhao, D. Asarnow, J. Biel, A. Bowen, J.R. Braxton, J. Chen, C.M. Chio, U.S. Chio, I. Deshpande, L. Doan, B. Faust, S. Flores, M. Jin, K. Kim, V.L. Lam, F. Li, J. Li, Y.-L. Li, Y. Li, X. Liu, M. Lo, K.E. Lopez, A.A. Melo, F.R. Moss, P. Nguyen, J. Paulino, K.I. Pawar, J.K. Peters, T.H. Pospiech, M. Safari, S. Sangwan, K. Schaefer, P.V. Thomas, A.C. Thwin, R. Trenker, E. Tse, T.K.M. Tsui, F. Wang, N. Whitis, Z. Yu, K. Zhang, Y. Zhang, F. Zhou, D. Saltzberg, Q.S.B. Consortium12†, A.J. Hodder, A.S. Shun-Shion, D.M. Williams, K.M. White, R. Rosales, T. Kehrer, L. Miorin, E. Moreno, A.H. Patel, S. Rihn, M.M. Khalid, A. Vallejo-Gracia, P. Fozouni, C.R. Simoneau, T.L. Roth, D. Wu, M.A. Karim, M. Ghoussaini, I. Dunham, F. Berardi, S. Weigang, M. Chazal, J. Park, J. Logue, M. McGrath, S. Weston, R. Haupt, C.J. Hastie, M. Elliott, F. Brown, K.A. Burness, E. Reid, M. Dorward, C. Johnson, S.G. Wilkinson, A. Geyer, D.M. Giesel, C. Baillie, S. Raggett, H. Leech, R. Toth, N. Goodman, K.C. Keough, A.L. Lind, Z. Consortium‡, R.J. Klesh, K.R. Hemphill, J. Carlson-Stevermer, J. Oki, K. Holden, T. Maures, K.S. Pollard, A. Sali, D.A. Agard, Y. Cheng, J.S. Fraser, A. Frost, N. Jura, T. Kortemme, A. Manglik, D.R. Southworth, R.M. Stroud, D.R. Alessi, P. Davies, M.B. Frieman, T. Ideker, C. Abate, N. Jouvenet, G. Kochs, B. Shoichet, M. Ott, M. Palmarini, K.M. Shokat, A. García-Sastrex, J.A. Rassen, R. Grosse, O.S. Rosenberg, K.A. Verba, C.F. Basler, M. Vignuzzi, A.A. Peden, P. Beltrao, and N.J. Krogan. 2020. Comparative host-coronavirus protein interaction networks reveal pan-viral disease mechanisms. Science.

5. V’kovski, P., A. Kratzel, S. Steiner, H. Stalder, and V. Thiel. 2021. Coronavirus biology and replication: implications for SARS-CoV-2. Nat. Rev. Microbiol. 19:155–170.

6. von Brunn, A., C. Teepe, J.C. Simpson, R. Pepperkok, C.C. Friedel, R. Zimmer, R. Roberts, R. Baric, and J. Haas. 2007. Analysis of intraviral protein-protein interactions of the SARS coronavirus ORFeome. PloS One2:e459.

7. Benvenuto, D., S. Angeletti, M. Giovanetti, M. Bianchi, S. Pascarella, R. Cauda, M. Ciccozzi, and A. Cassone. 2020. Evolutionary analysis of SARS-CoV-2: how mutation of Non-Structural Protein 6 (NSP6) could affect viral autophagy. J. Infect. 81:e24–e27.

8. Cottam, E.M., H.J. Maier, M. Manifava, L.C. Vaux, P. Chandra-Schoenfelder, W. Gerner, P. Britton, N.T. Ktistakis, and T. Wileman. 2011. Coronavirus nsp6 proteins generate autophagosomes from the endoplasmic reticulum via an omegasome intermediate. Autophagy7:1335–1347.

9. Oostra, M., M.C. Hagemeijer, M. van Gent, C.P.J. Bekker, E.G. te Lintelo, P.J.M. Rottier, and C.A.M. de Haan. 2008. Topology and membrane anchoring of the coronavirus replication complex: not all hydrophobic domains of nsp3 and nsp6 are membrane spanning. J. Virol. 82:12392–12405.

10. Angelini, M.M., M. Akhlaghpour, B.W. Neuman, and M.J. Buchmeier. 2013. Severe acute respiratory syndrome coronavirus nonstructural proteins 3, 4, and 6 induce double-membrane vesicles. mBio. 4.

11. Schweizer, L., and L. Mueller. 2014. Chapter 7 - Protein Conformational Dynamics and Signaling in Evolution and Pathophysiology. In: Arey BJ, editor. Biased Signaling in Physiology, Pharmacology and Therapeutics. San Diego: Academic Press. pp. 209–249.

12. Uversky, V.N. 2009. Intrinsically Disordered Proteins and Their Environment: Effects of Strong Denaturants, Temperature, pH, Counter Ions, Membranes, Binding Partners, Osmolytes, and Macromolecular Crowding. Protein J. 28:305–325.

13. Oldfield, C.J., J. Meng, J.Y. Yang, M.Q. Yang, V.N. Uversky, and A.K. Dunker. 2008. Flexible nets: disorder and induced fit in the associations of p53 and 14-3-3 with their partners. BMC Genomics. 9 Suppl 1:S1.

14. Kumar, D., P.M. Mishra, K. Gadhave, and R. Giri. 2020. Conformational dynamics of p53 N-terminal TAD2 region under different solvent conditions. Arch. Biochem. Biophys. 689:108459.

15. Kumar, A., P. Kumar, S. Kumari, V.N. Uversky, and R. Giri. 2020. Folding and structural polymorphism of p53 C-terminal domain: One peptide with many conformations. Arch. Biochem. Biophys. 684:108342.

16. Bhardwaj, T., K.U. Saumya, P. Kumar, N. Sharma, K. Gadhave, V.N. Uversky, and R. Giri. Japanese encephalitis virus – exploring the dark proteome and disorder–function paradigm. FEBS J. n/a.

17. Borcherds, W., F.-X. Theillet, A. Katzer, A. Finzel, K.M. Mishall, A.T. Powell, H. Wu, W. Manieri, C. Dieterich, P. Selenko, A. Loewer, and G.W. Daughdrill. 2014. Disorder and residual helicity alter p53-Mdm2 binding affinity and signaling in cells. Nat. Chem. Biol. 10:1000–1002.

18. Ghosh, A., A.S. Pithadia, J. Bhat, S. Bera, A. Midya, C.A. Fierke, A. Ramamoorthy, and A. Bhunia. 2015. Self-Assembly of a 9-Residue Amyloid-Forming Peptide Fragment of SARS Corona Virus E-protein: Mechanism of Self Aggregation and Amyloid-Inhibition of hIAPP. Biochemistry54:2249–2261.

19. Whitesides, G.M., J.P. Mathias, and C.T. Seto. 1991. Molecular self-assembly and nanochemistry: a chemical strategy for the synthesis of nanostructures. Science254:1312– 1319.

20. Thévenet, P., Y. Shen, J. Maupetit, F. Guyon, P. Derreumaux, and P. Tufféry. 2012. PEP-FOLD: an updated de novo structure prediction server for both linear and disulfide bonded cyclic peptides. Nucleic Acids Res. 40:W288–293.

21. Shen, Y., J. Maupetit, P. Derreumaux, and P. Tufféry. 2014. Improved PEP-FOLD Approach for Peptide and Miniprotein Structure Prediction. J. Chem. Theory Comput. 10:4745–4758.

22. Källberg, M., G. Margaryan, S. Wang, J. Ma, and J. Xu. 2014. RaptorX server: a resource for template-based protein structure modeling. Methods Mol. Biol. Clifton NJ1137:17–27.

23. Pettersen, E.F., T.D. Goddard, C.C. Huang, G.S. Couch, D.M. Greenblatt, E.C. Meng, and T.E. Ferrin. 2004. UCSF Chimera--a visualization system for exploratory research and analysis. J. Comput. Chem. 25:1605–1612.

24. Lee, J., X. Cheng, J.M. Swails, M.S. Yeom, P.K. Eastman, J.A. Lemkul, S. Wei, J. Buckner, J.C. Jeong, Y. Qi, S. Jo, V.S. Pande, D.A. Case, C.L. Brooks, A.D. MacKerell, J.B. Klauda, and W. Im. 2016. CHARMM-GUI Input Generator for NAMD, GROMACS, AMBER, OpenMM, and CHARMM/OpenMM Simulations Using the CHARMM36 Additive Force Field. J. Chem. Theory Comput. 12:405–413.

25. Kumar, A., A. Kumar, P. Kumar, N. Garg, and R. Giri. 2021. SARS-CoV-2 NSP1 C-terminal (residues 131–180) is an intrinsically disordered region in isolation. Curr. Res. Virol. Sci. 2:100007.

26. Jephthah, S., L. Staby, B.B. Kragelund, and M. Skepö. 2019. Temperature Dependence of Intrinsically Disordered Proteins in Simulations: What are We Missing? J. Chem. Theory Comput. 15:2672–2683.

27. Kjaergaard, M., A.-B. Nørholm, R. Hendus□Altenburger, S.F. Pedersen, F.M. Poulsen, and B.B. Kragelund. 2010. Temperature-dependent structural changes in intrinsically disordered proteins: Formation of α□helices or loss of polyproline II? Protein Sci. 19:1555–1564.

