## Supplementary figure 1, Supplementary figure 2, Supplementary figure 3 for "Investigating the conformational dynamics of SARS-CoV-2 NSP6 protein with emphasis on non-transmembrane 91-112 & 231-290 regions"

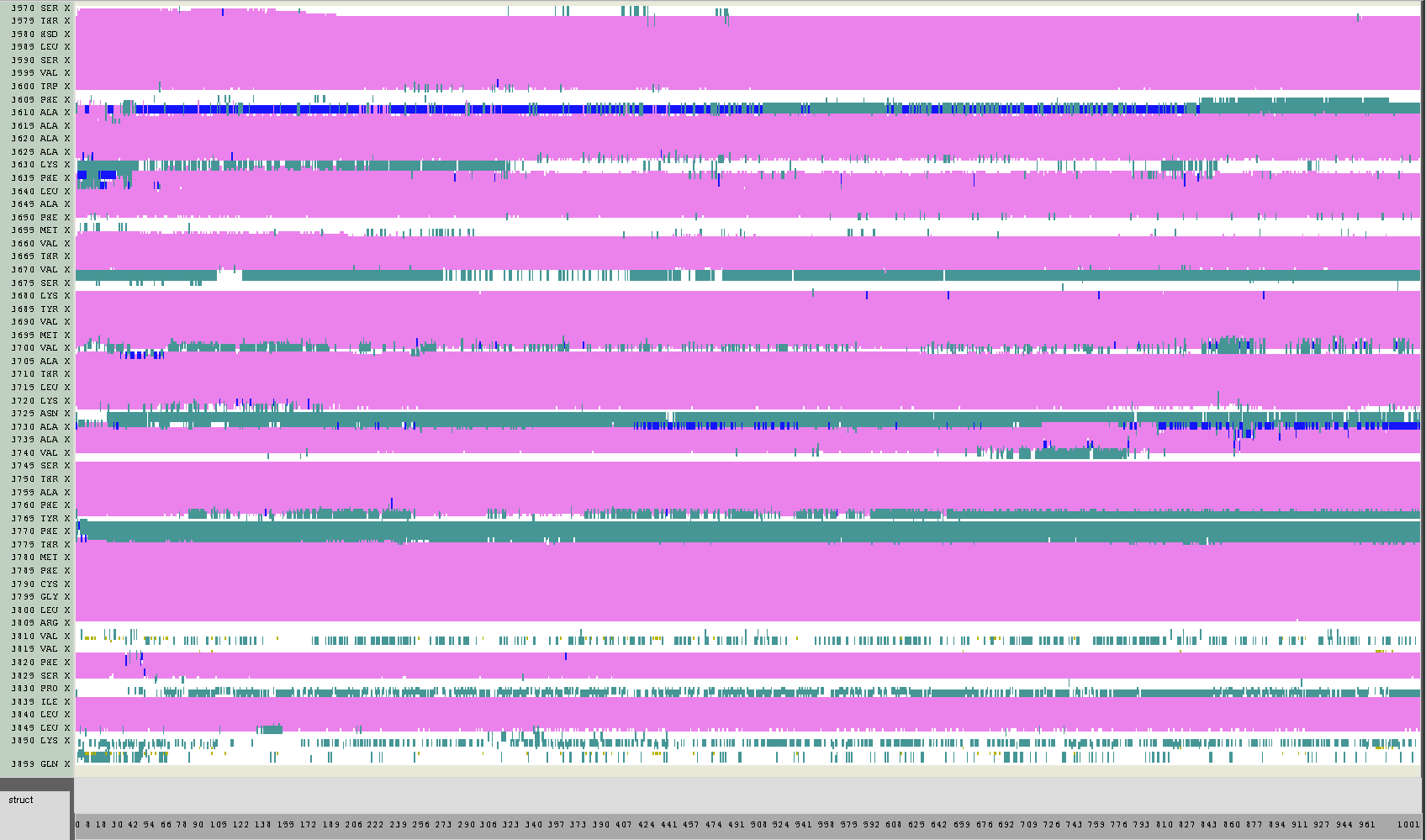

**Supplementary figure 1:** Secondary structure timeline representation of full-length NSP6 from molecular dynamics simulation trajectory up to 100ns using VMD Timeline extension. The alpha helical, 3-10 helix, and turns are shown with magenta, blue, and aqua colors, respectively.

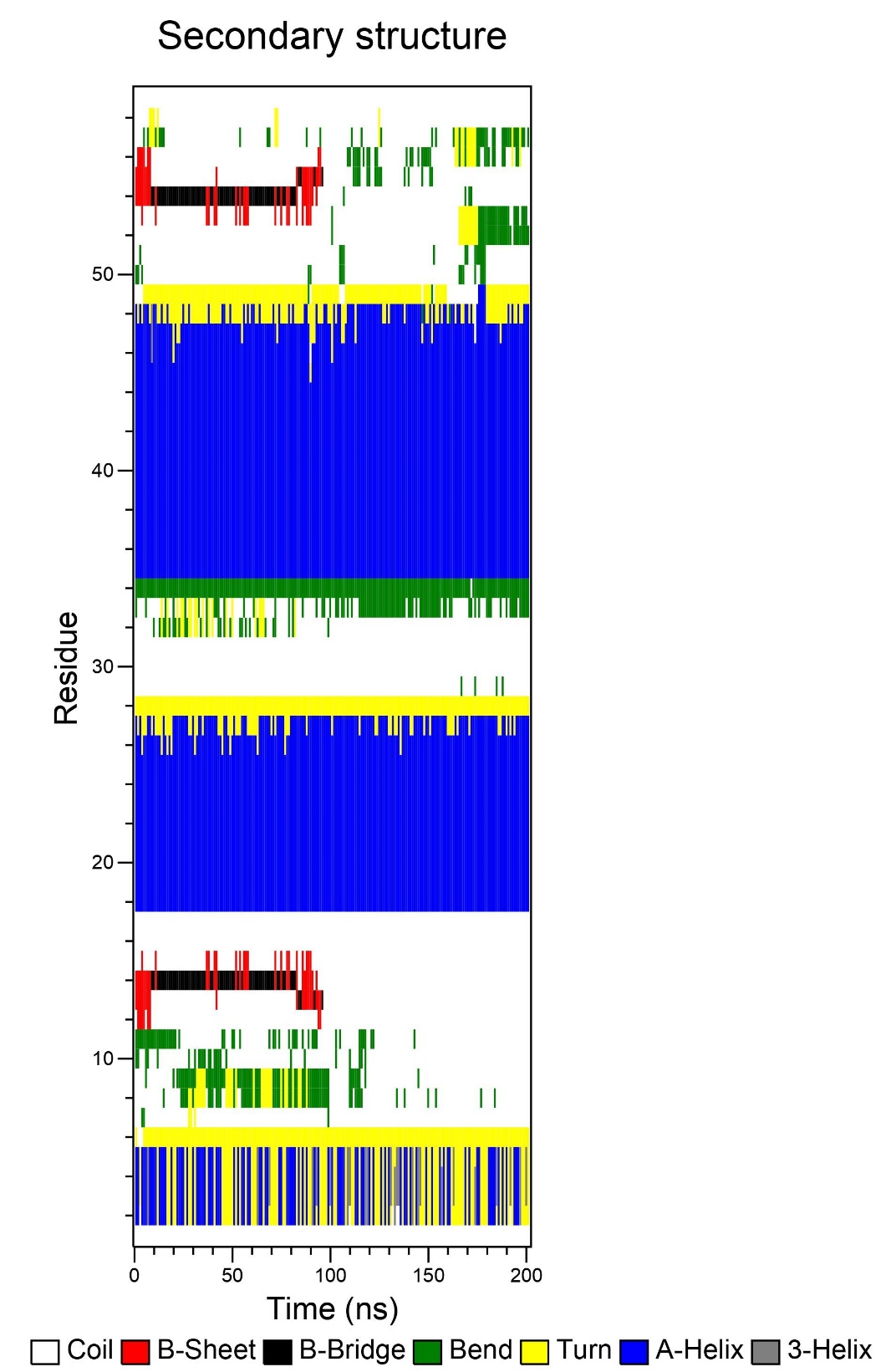

**Supplementary figure 2:** Secondary structure timeline representation of NSP6 C-terminal (residues 231-290) from molecular dynamics simulation trajectory up to 200ns using gromacs *do_dssp* command. The color schemes are shown in figure.

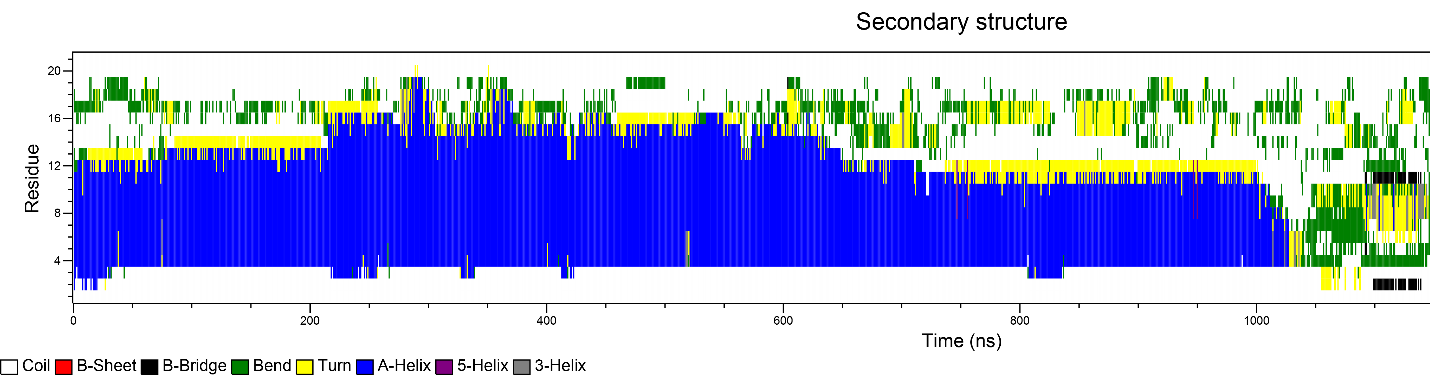

**Supplementary figure 3:** Secondary structure timeline representation of NSP6 non-transmembrane region (residues 91-112) from molecular dynamics simulation trajectory up to 1500ns (1.5µs) using gromacs *do_dssp* command. The color schemes are shown in figure.
